# Ecosystem models of Lake Victoria (East Africa): exploring the sensitivity of ecosystem effects of fishing to model choice

**DOI:** 10.1101/489260

**Authors:** Vianny Natugonza, Cameron Ainsworth, Erla Sturludóttir, Laban Musinguzi, Richard Ogutu-Ohwayo, Tumi Tomasson, Chrisphine Nyamweya, Gunnar Stefansson

## Abstract

Ecosystem simulation models are valuable tools for strengthening and promoting ecosystem-based fisheries management (EBFM). However, utility of these models in practical fisheries management is often undermined by lack of simple means to test the effect of uncertainty on model outputs. Recently, the use of multiple ecosystem models has been recommended as an ‘insurance’ against effects of uncertainty that comes with modelling complex systems. The assumption is that if models with different structure and formulation give consistent results, then, policy prescriptions are robust (i.e. less sensitive to model choice). However, information on the behaviour of trends from structurally-distinct ecosystem models with respect to changes in fishing conditions is limited, especially for freshwater systems. In this study, we compared outputs of two ecosystem models, Ecopath with Ecosim (EwE) and Atlantis, for Lake Victoria under different fishing pressure scenarios. We compared model behaviour at the ecosystem level, and also at a level of functional groups. At functional group level, we determined two questions: what is the change in the targeted group, and what are the consequent effects in other parts of the system? Overall results suggest that different model formulations can provide similar qualitative predictions (direction of change), especially for targeted groups with similar trophic interactions and adequate data for parameterization and calibration. However, considerable variations in predictions (where models predict opposite trends) may also occur due to inconsistencies in the strength of the aggregate multispecies interactions between species and models, and not necessarily due to model detail and complexity. Therefore, with more information and data, especially on diet, and comparable representation of feeding interactions across models, ecosystem models with distinct structure and formulation can give consistent policy evaluations for most biological groups.

## Introduction

### Ecosystem modelling for ecosystem-based fisheries management (EBMF)

In the recent years, calls for the implementation of ecosystem-based fisheries management (EBFM) have increased [1], despite the slow progress towards its adoption [2, 3]. The slow adoption of EBFM has largely been due to divergences in the interpretation among professionals [4, 5]. The advantages of EBFM are clearly understood. For example, it considers how fishing impacts entire ecosystem and fisheries through both direct and indirect mechanisms when formulating fisheries management strategies and actions [5].

Ecosystem simulation models can be used to evaluate ecosystem properties and provide information on the potential effects that changes in EBFM practices would have on the ecosystems [6]. Within the last two decades, ecosystem models have become popular tools for influencing and strengthening EBFM [7]. However, ecosystem models differ in detail of their biological processes and how they are represented, projection length and solution time steps [8]. This variation in model detail and assumptions introduces varying levels of uncertainty that often undermine utility of end-to-end models in practical fisheries management [9].

The high levels of uncertainty inherent in some ecosystem models means that no ecosystem model is perfect for all purposes under the EBFM framework [10]. This is exacerbated by the subjective nature of the modelling process as parameter estimation within the models is not possible. Although these models are constructed based on the knowledge of the system (i.e. to minimize process uncertainty), and also utilizing the best available data, these are not adequate safeguards to uncertainty that comes with modelling complex systems. In ecosystem models with intermediate complexity such as Ecopath with Ecosim (EwE), Monte Carlo algorithm is applied to examine the sensitivity of simulation results to the initial input parameters [11]. However, for complex end-to-end ecosystem models, such as Atlantis with thousands of parameters, full-scale sensitivity analysis is not feasible.

### The use of multiple ecosystem models

To limit on the effect of model uncertainty on policy recommendations, the use of multiple and complementary ecosystem models to provide input for management is strongly recommended [12, 13]. However, this requires a clear understanding of the level of robustness of results from different model formulations. Robustness here is considered to refer to consistency of performance across alternative model formulation, model uncertainty, and levels of perturbation intensity [14].

Multi-species models are multi-dimensional, and comparing them is generally a complex task. Consequently, recent investigations have focused on simpler approaches to understand how ecosystem impacts of fishing are sensitive to model choice using a range of indicators [15–19]. At the broadest level, these studies have found considerable coherence in general predictions (i.e. direction of change) across models but still with major differences observed for the multi-species effects. Whereas the general causes of discrepancies have been identified, including model structure and differences in representation of diets, some variations are ecosystem-specific [19].

The structural and functional differences between the multi-species models are huge. For example, EwE is a whole ecosystem biomass model, which is not spatially resolved unless coupled with Ecospace, where predation is regulated by explicit diet parameters and foraging vulnerability [11]. On the other hand, Atlantis is a whole ecosystem, age- and size-structured population model that is resolved in three dimensions with user-defined polygonal model zones and multiple depth layers [20, 21]. Predation in Atlantis is regulated by a diet preference matrix, but the actual resulting diet is subject to mouth-gape limitations and prey availability. The two modelling approaches have no systematic variation in assumptions; yet, they are designed almost to achieve the same ultimate goal: evaluation of system-level trade-offs of alternative management strategies. Determining whether the different model formulations predict similar outcomes in response to changes in fishing conditions is important in the EBFM context. Even where models predict different outcomes, such comparisons are useful in highlighting areas where different assumptions may lead to varying predictions, which can be used to improve the models.

### Ecosystem models of Lake Victoria (East Africa)

Considerable attempts have been made towards constructing ecosystem models for Lake Victoria to understand ecosystem dynamics (structure and functioning) as well as ecosystem-level effects of alternative fishery policies. Emphasis has been put on use of EwE and Atlantis modelling frameworks because of their popularity across the African Great Lakes [22], and generally across the globe [10, 23].

EwE and Atlantis models of Lake Victoria have been constructed to answer specific questions that are common to both models: food web structure and function and ecosystem effects of fishing [22]. However, to improve our confidence in results from these models, there is need for systematic analysis of sensitivity of ecosystem impacts of fishing to model structure and formulation.

In this paper, we compared the behaviour of EwE and Atlantis model simulations of the Lake Victoria ecosystem. We compared model behaviour at the ecosystem level, and also at a level of functional groups. The work described here is not intended to recommend one model over another. Rather, the main objective is to investigate how ecosystem effects of fishing are sensitive to model choice, and which ecosystem indicators are most sensitive to model uncertainty and complexity. Because the outputs of complex ecosystem models such as the Atlantis are huge, to ease comparisons, we aggregated the results and concentrated on comparing the behaviour of ecosystem indicators. For biomass-based indicators, results from Atlantis were aggregated to show trends through time, with no spatial and age-structure considerations.

## Materials and Methods

### Study area

Lake Victoria, located in East Africa (Fig 1), is the most productive freshwater lake in the world, with annual fish landings of about one million tonnes, and the second largest in terms of size (with a surface area of about 68,800 km^2^). The fishery currently employs more than one million people directly in fishing and other value-chain related activities; when their dependents are included, Lake Victoria supports local livelihoods of about four million people [24].

**Fig 1.**
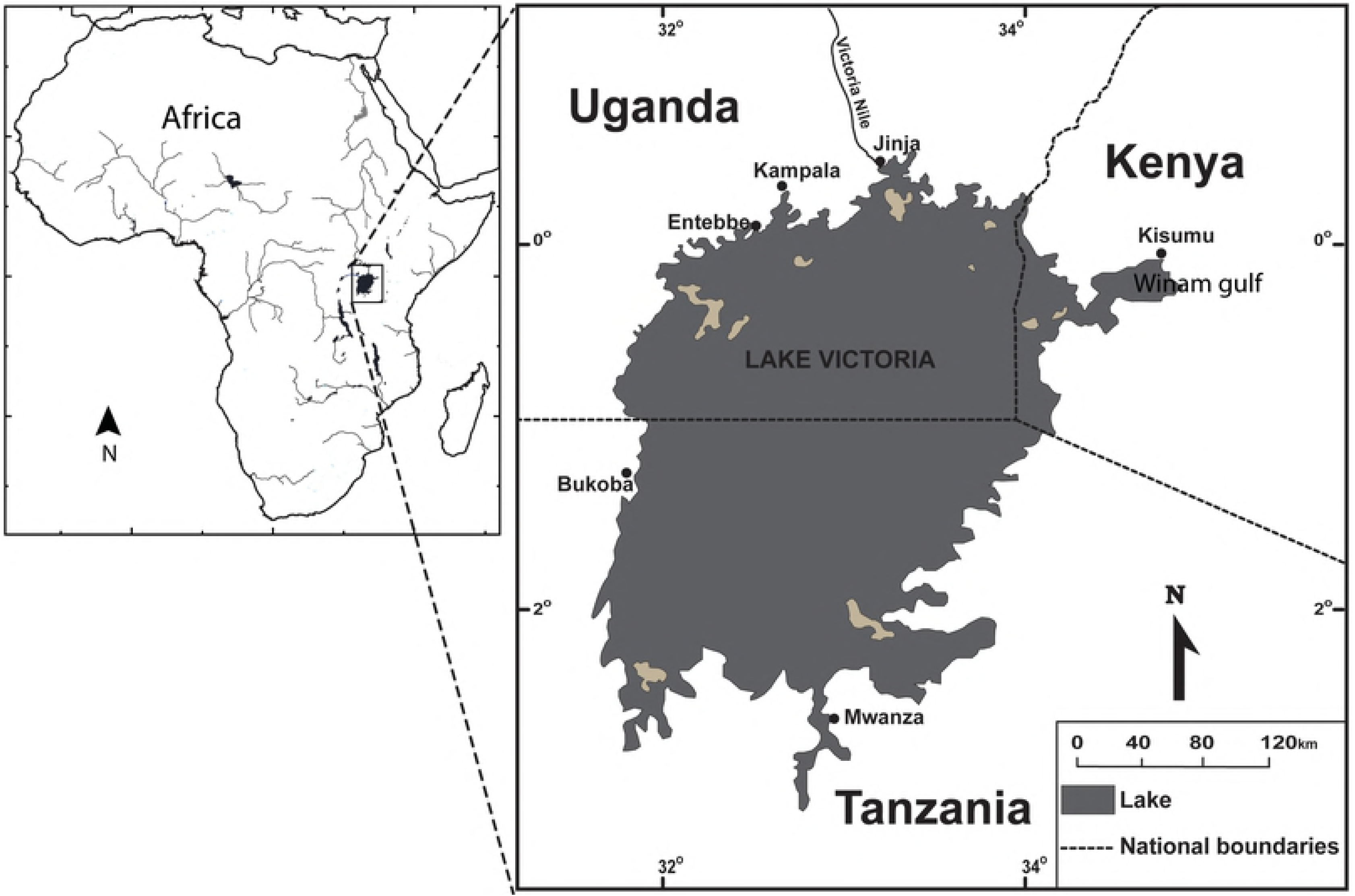
Location of Lake Victoria (East Africa) within Africa.

The present-day Lake Victoria fishery represents a massive transformation from the traditional and highly species-diverse fishery (i.e. before 1960s), known for its 500+ species of haplochromines, to a less species-diverse but highly productive and lucrative fishery dominated by introduced species especially Nile perch (*Lates niloticus*). An elaborate account of changes that have occurred, and how the fishery has persisted amidst multiple stressors e.g. species introductions, fishing, habitat degradation, eutrophication, and climate variability and change can be found in published literature [25–27].

## Modelling frameworks

### Ecopath with Ecosim (EwE)

The EwE modelling suite has been widely documented [11]. Briefly, EwE has an ecosystem trophic mass balance routine (Ecopath), where an ecosystem is partitioned into functional groups based on ecological roles and feeding interactions. Biomass flows in an ecosystem are regulated by gains (consumption, production, and immigration) and losses (mortality and emigration), through predator-prey relationships. For each functional group, the net difference between gains and losses is equal to the instantaneous rate of biomass change, which is parameterized with Biomass Accumulation. Key model parameters include biomass per unit of habitat area, production rate per unit of biomass, consumption rate per unit of biomass of predator, and ecotrophic efficiency (EE, the proportion of production that is utilized in the system). The model uses the input data along with algorithms and a routine for matrix inversion to estimate one missing basic parameter for each functional group, usually the EE. The Trophic level (TL) of each functional group is calculated on the basis of average annual predation by aggregating diet data. Primary producers and detritus are assigned a TL of 1, and the TL of consumer groups is calculated as the biomass-weighted average TL of its prey +1.

The time dynamic routine of EwE, Ecosim, uses Ecopath parameters to provide predictions of biomass and catch rates of each group as affected directly by fishing, predation, and change in food availability, and indirectly by fishing or predation on other groups in the system. Predation is governed by the concept of foraging arena, where species are divided into vulnerable and non-vulnerable components, such that the overall feeding rate is somehow limited by prey density. Calibration is achieved by adjusting diet and vulnerabilities until satisfactory fits are achieved.

### Atlantis

The Atlantis modelling framework has also been described elsewhere [20, 21]. Briefly, Atlantis is a deterministic, spatially resolved tool that is based on dynamically coupled biophysical and fisheries sub models (consumption, biological production, waste production, reproduction, habitat dependency, age structure, mortality, decomposition, and microbial cycles). Biophysical and biological processes are modelled in interconnected cells representing major features of the physical environment. The spatial domain is resolved in three dimensions using irregular polygons defined by the modeller to represent biogeographic features. Exchange of biomass occurs between polygons according to seasonal migration and foraging behaviour, while water fluxes (which control advection of nutrients and plankton), heat, and salinity flux across boundaries are represented by a coupled hydrodynamic model.

Functional groups, as with EwE, are determined based on ecological roles, ontogenetic behaviour and feeding interactions, except that vertebrates in Atlantis are represented as age-structured groups and lower trophic groups as biomass pools. The flow of energy is tracked as nitrogen, which in all vertebrate groups is partitioned into structural and reserve nitrogen. Structural nitrogen determines growth, while reserve nitrogen (whose amount varies depending on the food intake) is used for reproduction. The model simulates dynamic feeding interactions, with all functional feeding responses based on a modified Holling type II response. Trophic levels of model groups are computed on the same basis as in EwE.

### Operating models

In this study, we used the EwE and Atlantis models constructed for Lake Victoria as operating models. The models were constructed with an ultimate goal of exploring the ecosystem impacts of fishing, making it possible to compare the model behaviour under various fishing pressure scenarios. Fig 2 shows a summarized representation of the major features for the two models used in this study. The two models are similar in spatial extent (3.05°S to 0.55°N and 31.5° to 34.88°E), covering the area of approximately 68,800 km^2^, and were constructed to represent the ecosystem of Lake Victoria during the period when most of non-native species had just been introduced i.e. 1958 for Atlantis and 1960 for EwE. The calibration approach in two models differs substantially, but the period is comparable.

**Fig 2.**
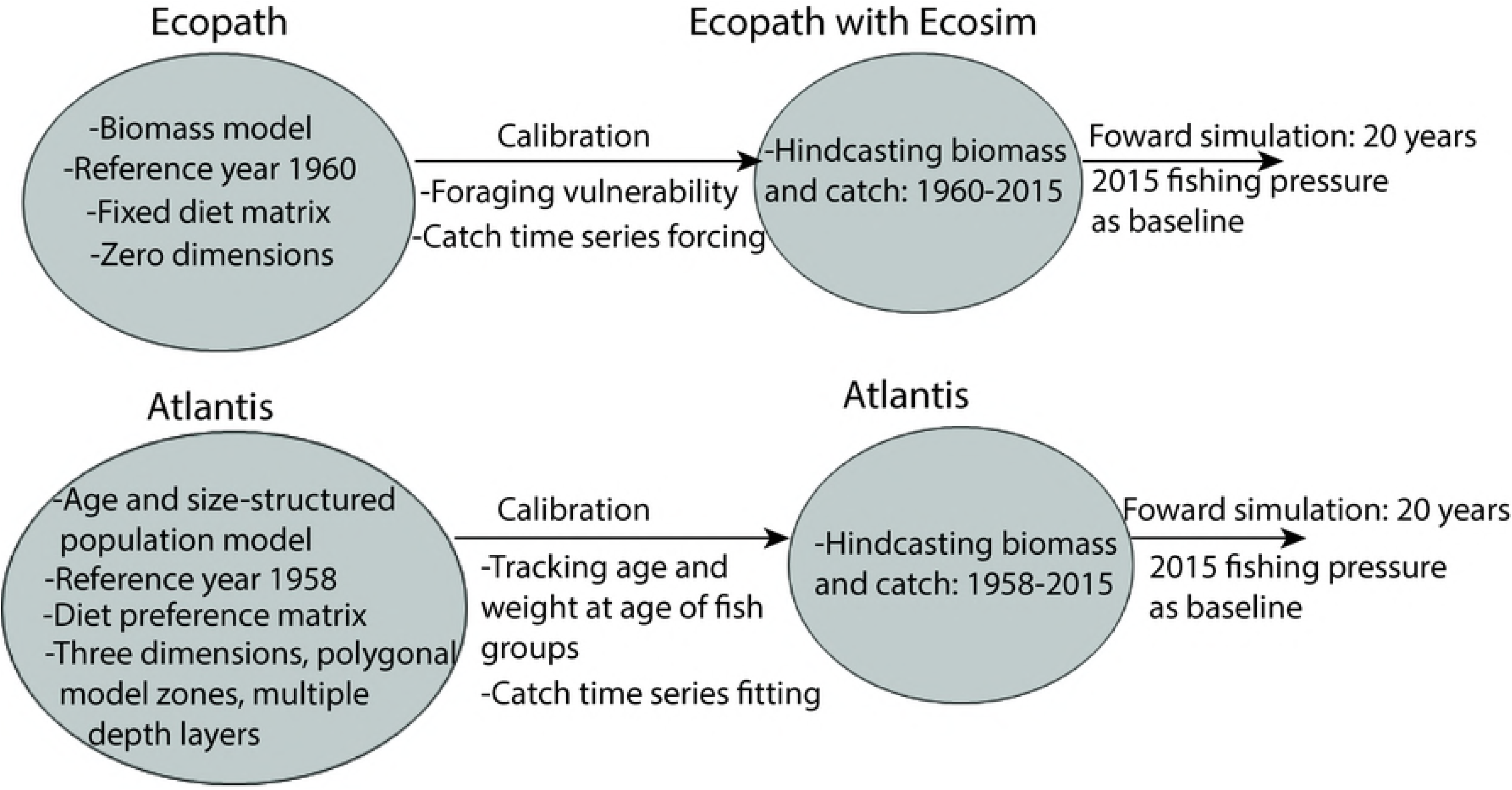
Schematic diagram showing the major features of EwE and Atlantis models for Lake Victoria used in this study.

The detailed EwE model used in this study (including set up, parameterization, and calibration) can be found at https://doi.org/10.6084/m9.figshare.7306820.v2. The model comprises 25 groups, including fish eating birds, the Nile crocodile, 15 fish groups (either as individual fish species or several species grouped together based on similarity in life history, habitat or diet), three invertebrate groups, two producer groups, and a detrital group (Table 1). Haplochromines, which is a group of major ecological importance (forage group), are modelled in one group, differing from Atlantis where haplochromines are modelled in three groups (Table 1). Nile perch, another group of focus in the fishing scenarios (see below), is also modelled as a single group, despite the species’ dietary preferences related to size [29]. Although Nile perch is also modelled as one in Atlantis model, it is divided into 10 age classes [31]; and therefore, the juvenile and adult individuals can have different diet and spatial distribution. In EwE, this is only modelled implicitly by including all possible prey for juvenile and adult Nile perch in the same diet matrix.

**Table 1.**
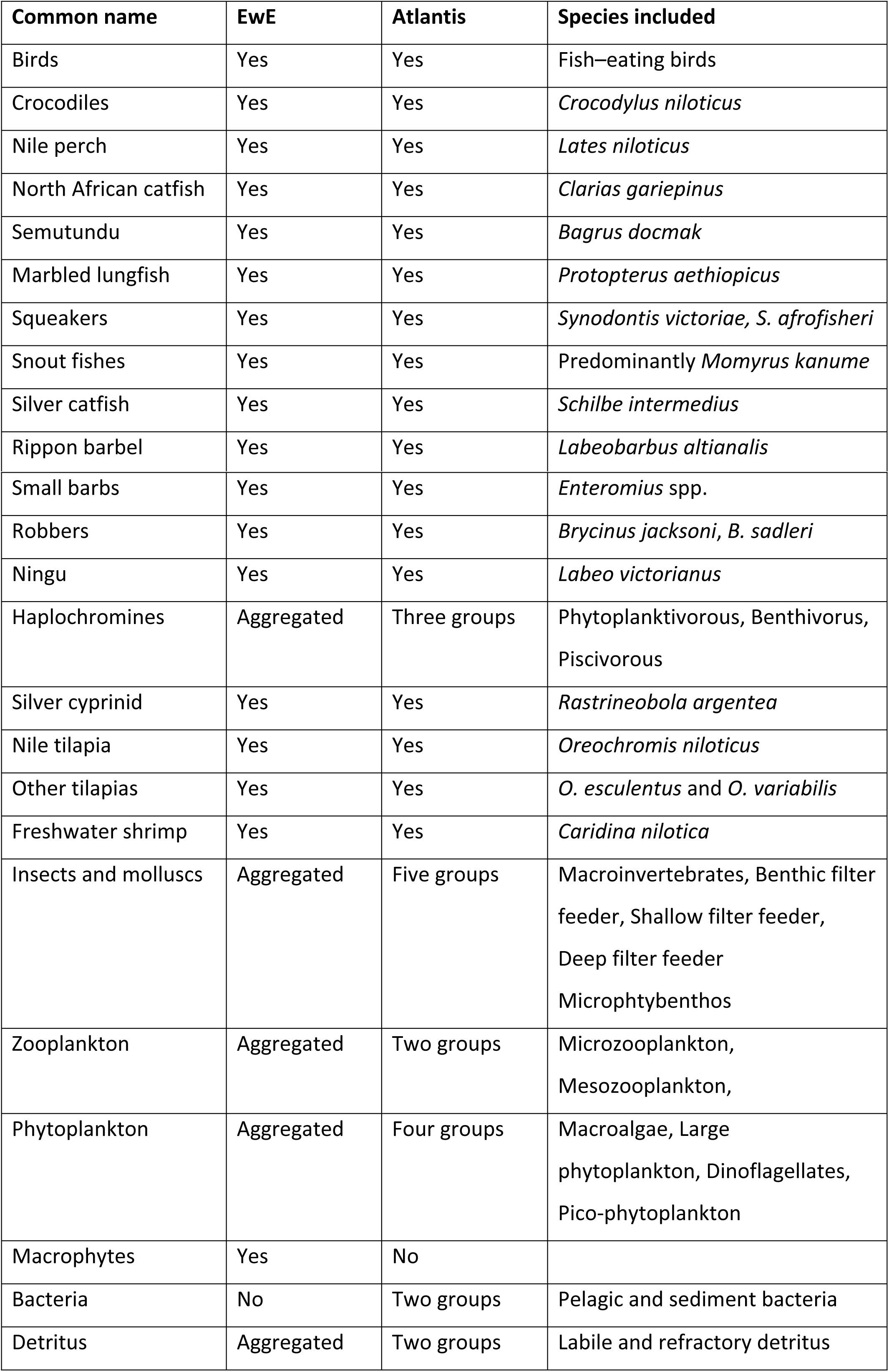
Functional groups used in the Lake Victoria EwE and Atlantis models.

The Atlantis model used in this study has been described in detail elsewhere [31, 32]. A complete set up of this model was retrieved from https://doi.org/10.6084/m9.figshare.4036077.v1. The model has 12 unique spatial regions, each region with 1-3 depth layers depending on the total depth, and a total of 34 of biological groups (i.e. 17 fish groups, fish eating birds, Nile crocodile, nine invertebrate and six primary producer groups). The 19 vertebrate groups are modelled as age-structured components, while the remaining 15 lower trophic groups are modelled as biomass pools.

Except for haplochromines, which are separated into three groups in Atlantis (Table 1), the choice of functional grouping at the vertebrate level for the two models is the same, although representation of diet is quite different (Fig 3). For the invertebrate and producer groups, the choice of functional groups differ substantially across models. Atlantis model has nine invertebrates groups and six producer groups compared to three invertebrate and two producer groups in the EwE model (Table 1). The detrital group in the Atlantis model is also divided into refractory and labile detritus. Therefore, our analysis focuses on groups that are comparable across models (Fig 3), excluding fish eating birds and crocodiles. For haplochromines, results for the three groups from Atlantis are aggregated and presented as one group.

**Fig 3.**
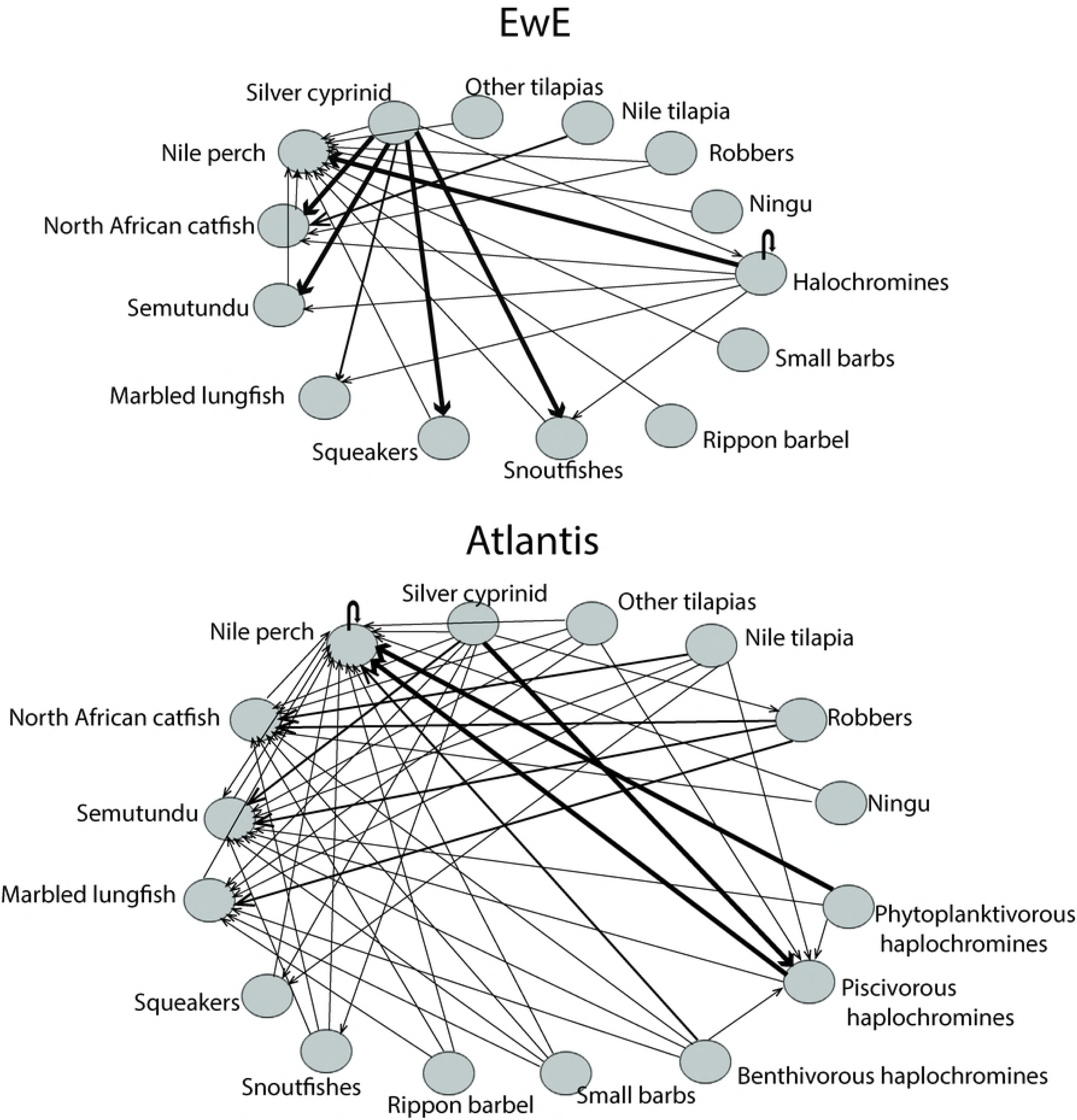
Schematic representation of predation interactions in EwE and Atlantis models of the Lake Victoria ecosystem. Model groups shown here are only for fish species, which are fairly represented in both models, to ease comparisons. Note that arrows move towards the predators and arrow thickness is consistent with the contribution of prey to the predator’s diet. Thick and black arrows indicate that the prey species makes up more than 30% of the predator’s diet, while thin arrows indicate that the prey species makes up less than 5% of the predator’s diet.

### Fishing scenarios

We focused on Nile perch and haplochromines in our fishing scenarios because of their greatest economic and ecological importance in the Lake Victoria ecosystem [33]. In addition, these groups are the most studied on the lake; we assume their representation in both models is fairly grounded in data, and their projections are less affected by data uncertainty compared to lesser-studied species. The fishing mortality for the last year of each historical model run (2015) was taken as the baseline fishing pressure. In the first and second scenarios, we reduced and increased, respectively, Nile perch fishing pressure by 40% from the baseline level. For the third scenario, we halted fishing of haplochromines (the major prey for Nile perch, see Fig 3). We also included the status quo scenario, where we maintained fishing pressure for all functional groups at the baseline level (i.e. as of 2015). We included the status quo scenario because the ecosystem would be expected to change under any level of fishing, and therefore the final results of the status quo scenario may not necessarily be the same as baseline values. For each scenario, biomass and catch for the individual species/groups were projected for 20 years into the future, and results are presented at the end of the projection period relative to the baseline (2015) values.

### Ecosystem indicators for comparison

Ecosystem indicators spanning a wide range of processes and biological groups have been used in several studies to detect a range of impacts from fishing [14]. To compare the changes that occur at a species/group level in response to fishing pressure scenarios, we looked at biomass of individual groups for each model but focused only on fish groups as they were represented in both models. We calculated Pearson correlation coefficient (*r*) for every functional group to examine the consistency of trends from both models under each fishing scenario. Our focus was on the direction of change in relative projections; so our subsequent interpretation of we use the term “consistency” to refer to any positive value of *r* and “inconsistency” to refer to negative values *r*.

Community-level indicators, on the other hand, are useful for detecting ecosystem-level changes [14]. These include relative abundance of key functional groups (e.g. piscivores and planktivores), mean TL in community and catch. Aggregating model groups into feeding guilds of fish species with broadly similar diets i.e. piscivores and planktivors is important because these feeding guilds are expected to respond to fishing pressure more predictably than individual species [28]. For instance, relative biomasses of piscivores and planktivores can indicate a change in the trophic structure of the system, as can shift in TL of the catch. Functional groups in the piscivorore guild included Nile perch, North African catfish, Semutundu, Silver catfish, and piscivorus haplochromines (TL>3.0). The planktivore guild included groups such as Silver cyprinid, Nile tilapia, other tilapia, Robbers, Ningu, Small barbs, phytoplanktivorous/Benthivorous haplochromines. Since the haplochromines in EwE are not segregated, we used relative abundance of Lake Victoria’s haplochromine trophic guilds [34] to assign biomass to each group.

We calculated Mean TL in community (*MTL_biomass_*) as the average TL of the model groups, weighted by their biomass according to equation 1

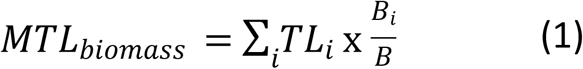

where *TL_i_* and *B_i_* are the trophic level and biomass of model group *i*, and *B* is the total biomass of all the fish groups (see Table 1). We only considered fish groups to avoid the influence of lower trophic planktonic groups (zooplankton and phytoplankton) that have comparatively greater biomasses. We preferred this approach because all the planktonic groups are not represented in all the models; therefore, focusing only on fish groups keeps the analysis comparable. Besides, the biomasses of planktonic groups can vary greatly with environmental effects, and such fluctuations may not be relevant to fisheries management.

We also calculated mean TL in catch (*MTL_catch_*) using the same approach as with MTL_biomass_, but using the biomass of catch for each model group rather than stock biomass i.e. as the mean TL of all landed fish, weighted by the biomass of catch (equation 2).

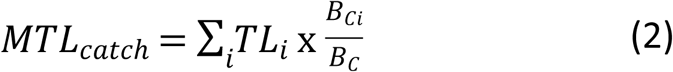

where *B_Ci_* is the biomass of catch of model group *i*. This indicator is important as it can signal to the depletion of high-trophic-level species i.e. ‘fishing down the food web’ [30].

## Results

### Biomasses of individual model groups

Fig 4 shows correlation values representing the change of trend of relative biomass of functional groups under different fishing pressure scenarios. Qualitative similarities (change in the same direction) between the two models are shown by functional groups with positive correlation values. Overall, the response to shifts in fishing pressure scenarios for individual functional groups was diverse across models, depending on the fishing pressure scenario in question. Projections with similar trends were observed for targeted groups and their prey/predator (depending on the strength of the feeding interaction), but large discrepancies were also observed especially for the indirect effects of the fishing pressure scenarios on non-target ‘distant’ groups. Only two groups (Nile perch and Nile tilapia) showed similar biomass trajectories (consistent trends) simultaneously in all the four scenarios.

**Fig 4.**
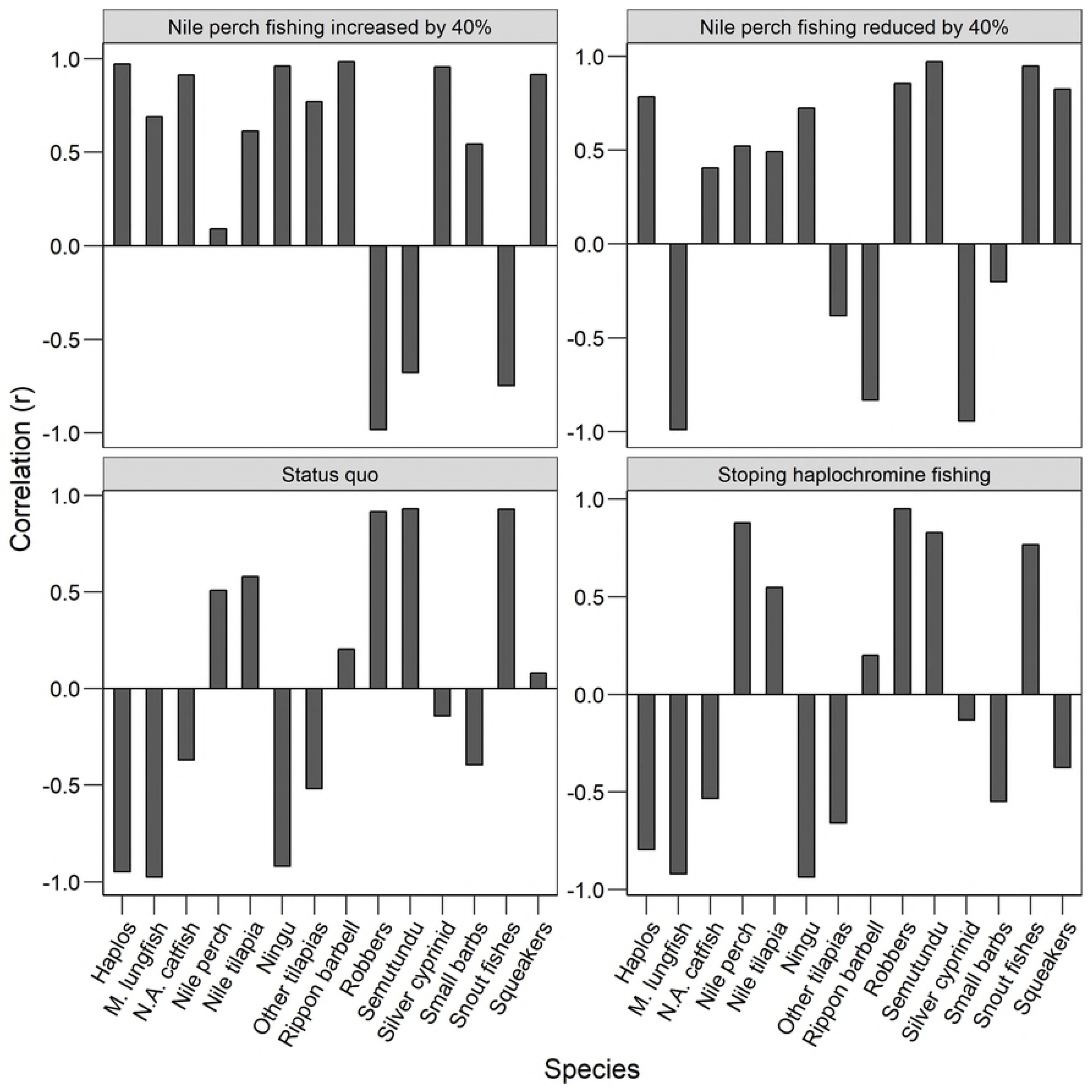
Correlation between the relative biomasses of species/groups projected by the two models under four different fishing scenarios. Haplos stands for haplochromines, M. lungfish is Marbled lungfish, and N.A. catfish is North African catfish.

The scenario of increasing Nile perch fishing pressure by 40% from baseline showed the highest level of consistency in biomass projections among functional groups (i.e. 11 out of the 14 model groups showed similar trends across models). The three groups whose trends differed were Robbers, Semutundu and snout fishes, where relative biomass increased in EwE but decreased in Atlantis. However, when Nile perch fishing pressure was instead reduced by 40% from the baseline, the number of groups with similar trends across models reduced to nine, although Robbers, Semutundu and snout fishes showed similar trends under this scenario. Only six groups (Nile perch, haplochromines, North African catfish, Nile tilapia, Ningu, and squeakers) showed similar direction of change across models under the two contrasting Nile perch fishing pressure scenarios.

The scenario of halting haplochromine fishing yielded the least number of groups with similar direction of change in biomass (i.e. six out of the 14 model groups). Unexpectedly, the response of haplochromines was also inconsistent, although the response of its major predator, Nile perch, was consistent under this scenario. The response of individual model groups under this fishing scenario was quintessentially similar to the status quo scenario. With the exception other tilapias, where Atlantis and EwE predicted an increase and decrease, respectively, the rest of the groups (Marbled lungfish, North African catfish, Ningu, Silver cyprinid, small barbs) decreased in Atlantis but increased in EwE.

Fig 5 shows change in predicted biomass by the two models under fishing pressure scenarios at the end of the simulation, relative to baseline. All outcomes of fishing pressure scenarios are compared at the end of 20 years, where values of zero indicate no change in biomass (relative to baseline levels). Qualitative agreements between models are shown by predictions in the same direction, indicated by bars on the same side of the zero line (either positive or negative sign). Quantitative agreements between models are shown by predictions with similar magnitude, indicated by bars with the same height. Generally, qualitative agreements were higher for the target groups (e.g. Nile perch, Nile tilapia, haplochromines, Silver cyprinid, Semutundu, and snoutfishes) than the non-target groups, although the magnitude of predictions differed substantially. Except for the scenario where Nile perch fishing was increased by 40% from baseline, Atlantis was generally more responsive to shifts in fishing pressure than EwE.

**Fig 5.**
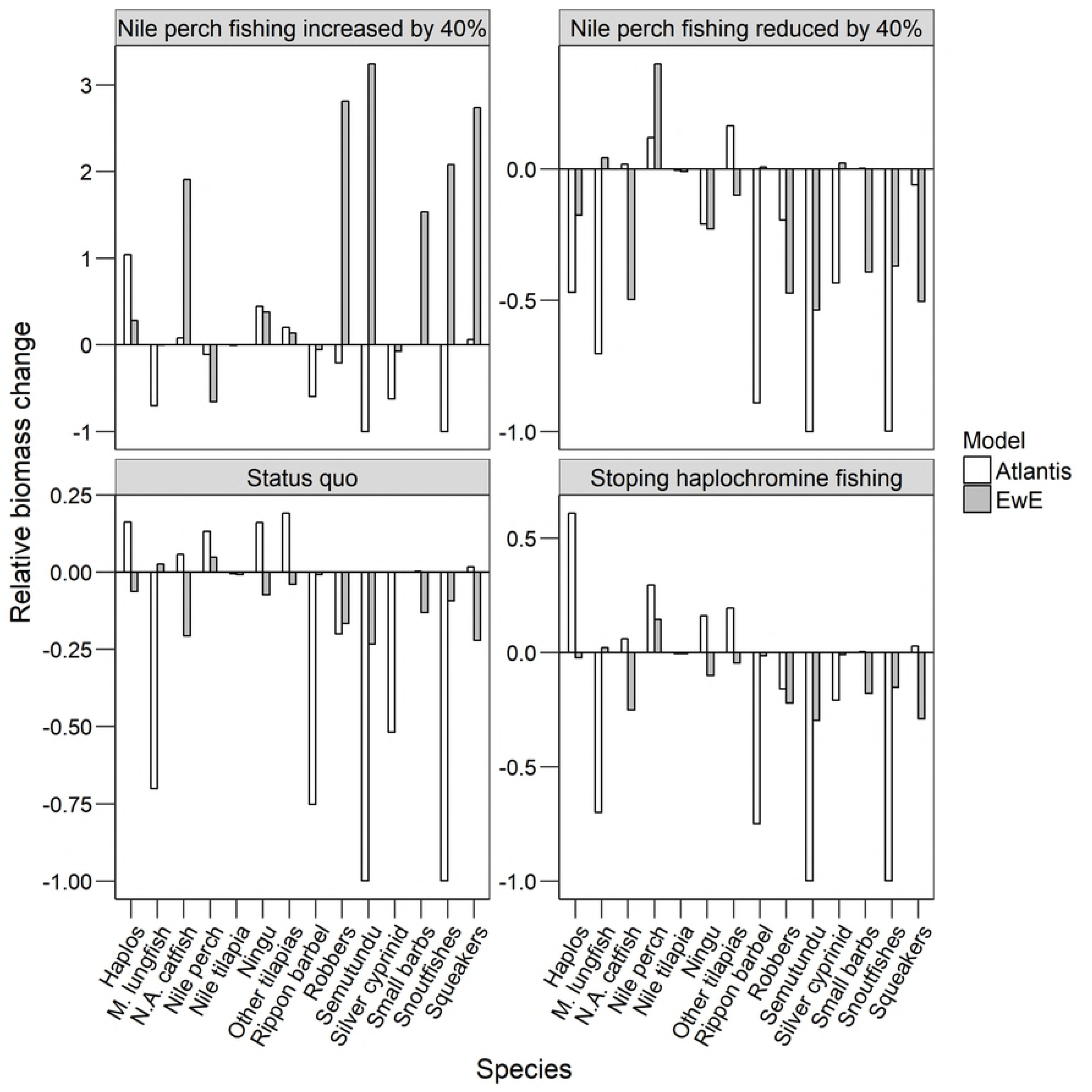
Relative change in biomass of functional groups at the end of forecasting period as predicted by Atlantis and EwE models.

Under the scenario of increasing Nile perch fishing pressure, Nile perch decreased both in EwE and Atlantis; however, the magnitude of the decrease was six times higher in EwE than Atlantis. As expected, the major prey for Nile perch (haplochromines) increased in both models, although Atlantis was more responsive than EwE. The response in other groups, except for Ningu and other tilapias, was highly variable, with EwE predicting an increase in biomass of most groups and Atlantis predicting a decrease.

Under the scenario of decreasing Nile perch fishing pressure, Nile perch increased while haplochromines decreased in both models, although the magnitude of decrease for haplochromines was higher (47%) in Atlantis than EwE (20%). For the rest of the groups, apart from Marbled lungfish and other tilapias, whose biomasses respectively increased and decreased in EwE and Atlantis (by at least 3%), the biomasses of other groups decreased in both models.

In the two other scenarios (maintaining status quo and halting haplochromine fishing), the predicted biomasses at the end of the simulation were highly variable across models, except for Nile perch, whose biomass increased, and three other groups (Snoutfishes, Semutundu, and Robbers) whose biomasses decreased consistently across models. Under these two scenarios, the responsiveness of the two models to shifts in fishing pressure was clearly higher in Atlantis than EwE.

### Ecosystem-level indicators

Fig 6 shows the proportional change in system-level indicators across the models under at the end each fishing pressure scenario. All indicators are shown as relative change from 2015 to 20135 for each scenario, where zero indicates no difference. Overall, ecosystem-level indicators were more consistent across models compared to the individual biomass-based indicators.

**Fig 6.**
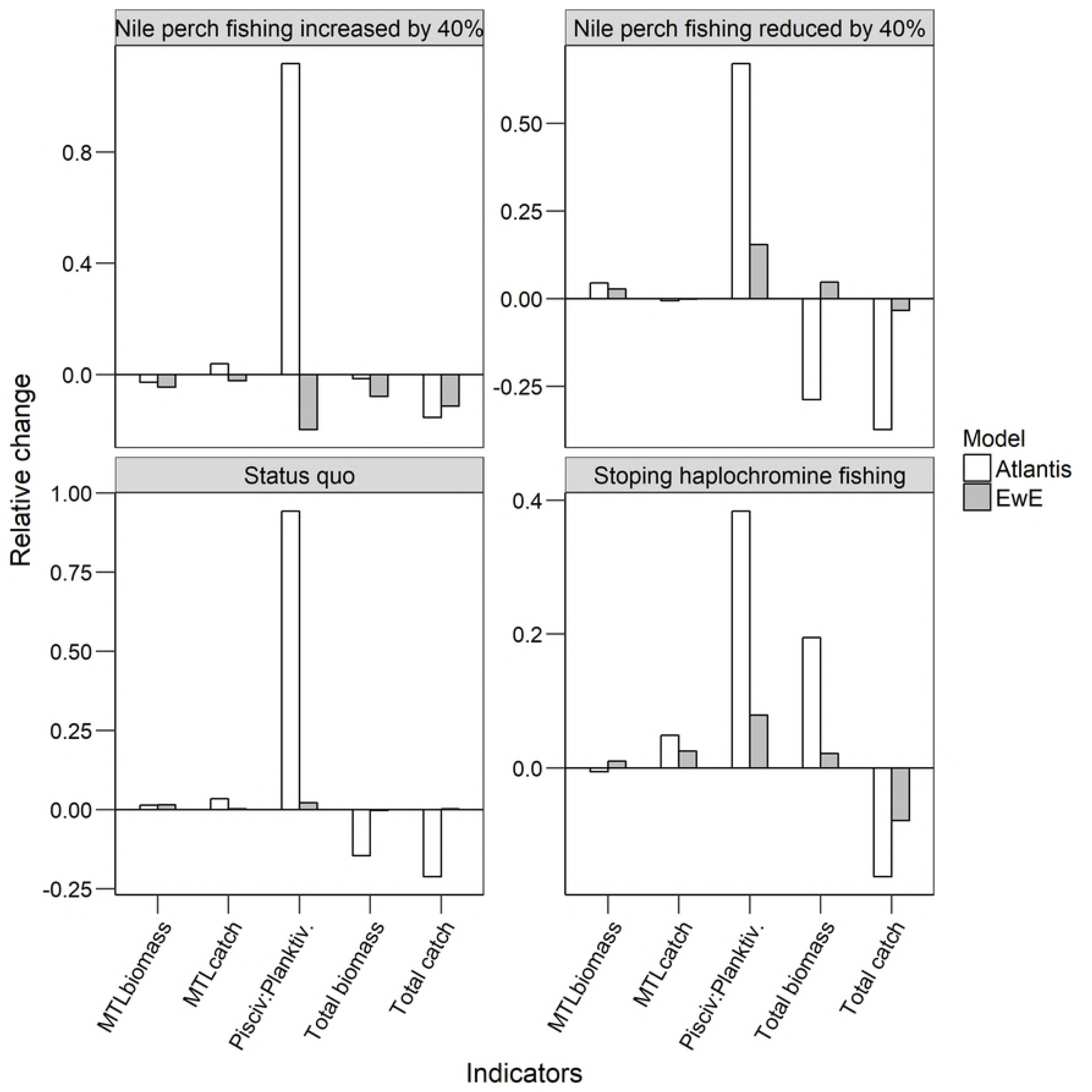
Relative change in system-level indicators in EwE and Atlantis under the four different fishing pressure scenarios. pisciv:planktiv stands for piscivorous to planktivorous ratio, MTL_biomass_ and MTL_catch_ are mean trophic level in community and catch, respectively.;

The biomass of piscivore guild relative to planktivore guild increased across models, except for the scenario of increasing Nile perch fishing, where Atlantis predicted a massive increase and EwE predicted a decrease. However, overall fish biomass decreased consistently across models.

MTL_biomass_ showed consistent direction of change across the models except under the scenario of halting haplochromine fishing, where the indicator value increased in Atlantis and decreased in EwE. MTL_biomass_ increased with the reduction Nile perch fishing and maintaining status quo, but decreased with an increase in Nile perch fishing. Similarly, MTL_catch_ was consistent across models, except the scenario of increasing Nile perch fishing pressure where Atlantis predicted an increase and EwE predicted a decrease. For the remaining scenarios, MTL_catch_ increased either by halting haplochromine fishing or maintaining status quo, but decreased by reducing Nile perch fishing pressure.

## Discussion

### Biomass of individual model groups

Ecosystem models are predominantly used to gain understanding of ecosystem-level processes and (in most cases) to indicate qualitative trends associated with changes in fishing (or some other form of forcing) conditions. Studies exploring consistency of ecosystem effects of fishing across models that have already taken place indicate that consistent general predictions (in terms of direction of change) can emerge from different model formulations, although considerable variations may occur in detailed model results especially for multispecies effects [14–19]. This is consistent with our general findings from this study. Our results suggest that the direction of change in biomass predictions is driven by trophic interactions, while the magnitude of change in predicted biomass depends on both the processes included in the model (model detail and complexity) as well as the strengths of feeding interactions.

The choice of biological groupings and representation of diets can greatly influence the level of connectivity between groups. This in turn has an effect on the projected magnitude of one species’ biomass or catch affected by other species’ fishing mortality. In our study, the effect of feeding interactions is illustrated by the two Nile perch fishing scenarios. Fig. 3 shows that the greatest proportion of Nile perch diet in both models is contributed by haplochromines. Reducing fishing pressure on Nile perch causes an expected increase in the abundance of Nile perch, which subsequently causes a decline in their preferred prey (haplochromines). The reverse is true as well owing to high fishing pressure and predation release on Nile perch and haplochromines, respectively. Although Nile perch feeds on other fishes such as Mabbled lungfish, Ningu, North African catfish, other tilapias, Robbers, Semutundu and squeakers, which all showed wide discrepancies in predicted biomass across models; these are weak feeding interactions, where each group contributes less than 3% in Nile perch diet. However, one striking feature about these groups (Mabbled lungfish, Ningu, North African catfish, other tilapias, Robbers, Semutundu and squeakers) is that they are all bentho-pelagic, largely feeding on invertebrates (not shown in Fig. 3) at the bottom sediment. Given that these groups don’t constitute a significant prey at the top of the food chain, changes in their abundance are governed by abundance of the lower TL invertebrate groups, whose grouping differs considerably across models (Table 1). Atlantis has nine invertebrate groups, while EwE has only three, with different feeding connections to high TL dependant groups. The discrepancies in biomass trend for these groups that depend on invertebrate prey can therefore be attributed to the differences in choice of functional groups at the bottom of the food chain, and not necessarily differences in model processes. This is especially true considering that Atlantis predicts a uniform decline in these groups under every fishing scenario.

Whereas the direction of change in model forecasts is largely governed by feeding interactions, model sensitivity to perturbation and the resulting magnitude of change in individual group biomasses seem to be driven both by the modelled processes and strength of the feeding dependencies. Studies that have previously compared Atlantis and EwE have found Atlantis to be less sensitive to changes in fishing pressure compared to EwE [16–18, 35, 36]. The authors have attributed the lower responsiveness of Atlantis to flexibility in feeding and incorporation of age structure and reproductive behaviour, which can delay the reproductive response of the population. In Atlantis predation is regulated by a diet preference matrix, although the actual resulting diet a function of mouth-gape and prey availability, while predation in EwE is regulated by a fixed diet matrix and foraging vulnerability. Fig 3 shows Atlantis model of Lake Victoria with more feeding linkages amongst compartments than EwE. This feeding flexibility in the Atlantis model, in addition to the ‘delaying’ model processes, were expected to dampen the sensitivity of predators to shifts in abundance of prey and result into lower responsiveness of Atlantis than EwE. However, this only occurred for Nile perch under the scenario where Nile perch fishing pressure was reduced; the magnitude of change for Nile perch in Atlantis was lower than in EwE (Fig. 5). For the rest of the groups, Atlantis was largely more sensitive to fishing than EwE, despite incorporating the delaying features of age structure and reproductive behaviour as well as allowing for diet flexibility. In this case, the strengths of diet dependencies likely outweighed the delaying system features.

### Ecosystem-level indicators

The shifts in biomass-weighted TL in community and catch, especially for the two contrasting Nile perch scenarios, were all consistent with expectation. Nile perch is a voracious predator at the top of the food chain; intensifying exploitation of this group decrease the average TL of the community, and the reverse occurs when Nile perch increases following a reduction in fishing. However, the sensitivity of the indicator is very low, which is caused by the large biomass of Silver cyprinid (a high-biomass pelagic zooplanktivore) that dampens the change in MTL_biomass_.

Under the ‘fishing down’ the food web hypothesis [30], TL of catch is expected to decline in response to fishing due to the preferential depletion of high-trophic-level species. In EwE, the direction of change of this indicator with respect to increased exploitation of Nile perch (top predator) was consistent with the ‘fishing down’ hypothesis. In Atlantis, however, TL of catch increased with increasing fishing pressure on Nile perch. Whereas this seems counter-intuitive, it is not entirely surprising because the increase in catches of the predator in the short-term can increase TL of the catch, which seems to be the case with the scenario of increasing exploitation on Nile perch.

By examining the feeding guilds, we expected to observe a fishing-driven decline in the piscivore guild under the scenario of increased fishing pressure on Nile perch. In turn, we expected this to cause an increase in the planktivore guild, which are major prey for the piscivore guild. Whereas results of EwE were consistent with this expectation, Atlantis predicted the opposite. The piscivorous to planktivorus ratio increased (substantially) in Atlantis even under heavy exploitation of Nile perch. This can be attributed to the rapid decline in Silver cyprinid, a dominant pelagic planktivore, possibly due to competition with haplochromines following predation release from intensively fished Nile perch. The rapid decline of Silver cyprinid cancels out any effect of small decline in Nile perch because when this indicator is calculated without the Silver cyprinid under the same scenario, the results are consistent with the above expectation.

## Conclusions

The overall model structure and formulation can provide similar qualitative predictions (direction of change), especially for groups with similar trophic interactions, although considerable variations may arise due to the differences in the strength of the aggregate multispecies interactions between species and models. Whereas qualitative model results depend on feeding interactions, model sensitivity to perturbation and the resulting magnitude of change in individual group biomasses are driven both by modelled processes and strength of the feeding dependencies. Availability of data for model parameterization and calibration also plays a role in the consistency of results across models. For example, Nile perch, Nile tilapia, and haplochromines (whose qualitative trends across models were all consistent in the scenarios tested) have been widely studied and documented, given their ecological and economic importance. The attention given to these species means that they are less likely to be affected by data uncertainty compared to lesser-studied species. Therefore, with more information and data, and comparable representation of trophic interactions across models, ecosystem models with distinct structure and formulation can easily give consistent policy evaluations for most of biological groups.

In the Lake Victoria Atlantis model, the strengths of diet dependencies exert bigger influence on model outcomes than any of the ‘delaying’ ecosystem features, such as age- and size structure or reproductive behaviour, which are common to Atlantis models. This is in regard to the higher sensitivity of Atlantis model to fishing pressure scenarios than EwE. Therefore, confidence in results from multiple models can be greatly enhanced by improving the accuracy of diet data through rigorous diet studies, especially for the less studied groups, and accurate definition of biological groups across models.

Ecosystem-level indicators are less sensitive to model choice compared to biomass of individual model groups; therefore, the actual ecosystem impacts of fishing from changes in these aggregated indicators needs to be interpreted with caution. This is true especially where the magnitude of change in indicator is small, as seen in this study, which could arise from opposite trends in several biological groups cancelling each other. Biomass information at the species level is still important for interpreting dynamics in ecosystem response to fishing. Even where models seem to give diverging results, this evaluation provides an account of possible changes from reference state and points to areas where different model considerations may lead to varying predictions, which can be used to improve the models.

## Acknowledgement

This work was supported by the United Nations University - Fisheries Training Program (UNU-FTP), Reykjavic, Iceland.

